# Comp2GPR: A Sequence-Driven Framework for Gene–Protein–Reaction Rule Reconstruction

**DOI:** 10.64898/2026.06.24.734174

**Authors:** Sandra Castillo

## Abstract

Accurate gene–protein–reaction (GPR) associations are essential for the predictive performance of genome-scale metabolic models (GEMs), as they define the mapping between genes, enzymes, and metabolic reactions. However, GPR rules are often incomplete or inconsistent due to limitations in annotation transfer and the ambiguous representation of multi-subunit protein complexes, leading to errors in downstream analyses such as gene essentiality prediction. Here, I introduce Comp2GPR, an automated pipeline for reconstructing GPR rules that integrates curated protein complex information with sequence-level evidence. Protein complexes were sourced from the Complex Portal and subjected to an AI-assisted curation workflow to retain only metabolically relevant assemblies. Comp2GPR combines deterministic sequence similarity mapping with explicit rule construction to generate Boolean GPR expressions that accurately represent obligate subunit relationships and isoenzyme redundancy. I evaluated the impact of the reconstructed GPR rules by integrating them into the Yeast9 metabolic model and comparing gene essentiality predictions with the original model. While global performance metrics remained largely unchanged, the updated model achieved a net improvement in prediction accuracy through gene-level corrections. Overall, Comp2GPR demonstrates that combining curated protein complex data with sequence-based validation improves the accuracy, interpretability, and reproducibility of GPR rules. The method provides a robust framework for enhancing metabolic model annotations and supports more reliable simulation-based analyses.

## Introduction

Genome-scale metabolic models (GEMs) have become essential tools for investigating cellular metabolism, enabling the systematic analysis of metabolic capabilities, prediction of gene essentiality, and rational strain engineering (13; 3; 20). A central component of these models is the set of gene–protein–reaction (GPR) associations, which define how genes map to enzymes and, ultimately, to metabolic reactions. These rules are typically represented as Boolean expressions encoding isoenzyme relationships (OR logic) and multi-subunit enzyme complexes (AND logic). Accurate GPR rules are therefore critical for ensuring that model simulations reflect the molecular constraints underlying metabolic function.

Despite their importance, the construction of GPR rules remains a major bottleneck in metabolic model development. In many reconstruction pipelines, GPR rules are absent, derived from annotation databases or transferred from existing models without explicit validation of protein complex composition or gene–protein mapping(7; 19; 15). This often leads to incomplete or incorrect representations of enzymatic machineries, particularly for multi-subunit complexes, where missing or incorrectly assigned components can introduce artificial bottlenecks or spurious alternative pathways.

As a consequence, inaccuracies in GPR rules propagate to downstream analyses, affecting flux predictions, gene essentiality assessments, and model-based design strategies.

Several approaches have been proposed to automate GPR rule reconstruction, including methods that integrate protein annotation data and curated databases to infer gene–reaction associations (5; 11). While these approaches improve scalability, they typically rely on indirect annotation-based mappings and database completeness. As a result, they may incorporate protein complexes that are not metabolically relevant or assign genes to complex components without sufficient sequence-level evidence, leading to ambiguous or over-permissive GPR rules. In particular, the accurate reconstruction of multi-subunit enzyme complexes remains a major challenge, as errors in AND relationships can significantly distort model behavior.

Recent evidence highlights the limitations of annotation-driven approaches, especially in complex reconstruction tasks. For example, methods relying primarily on annotation integration show markedly reduced accuracy when predicting GPR rules for multi-subunit enzyme complexes, indicating that database-driven inference alone is insufficient to capture obligate subunit relationships. These limitations underscore the need for approaches that directly validate gene–protein associations at the sequence level while ensuring that only metabolically relevant protein complexes are considered.

A key challenge in this context is the integration of protein complex knowledge into metabolic modeling. Databases such as Complex Portal provide extensive information on protein complex composition and function across organisms; however, they include a wide range of biological assemblies, many of which are unrelated to metabolism. Direct incorporation of such datasets therefore introduces noise and increases the risk of incorrect gene–reaction associations unless additional filtering and validation steps are applied.

In this work, I present **Comp2GPR**, an automated pipeline for reconstructing GPR rules that integrates curated protein complex information with sequence-level evidence. The approach combines an AI-assisted curation workflow to extract metabolically relevant complexes with a deterministic sequence similarity mapping strategy, enabling the construction of Boolean rules that explicitly represent obligate subunit relationships and isoenzyme redundancy. By enforcing one-to-one gene–protein assignments and filtering unsupported associations, Comp2GPR aims to improve the accuracy, interpretability, and reproducibility of GPR rules in genome-scale models.

I evaluated the performance of Comp2GPR by comparing reconstructed GPR rules to those of the Yeast9 consensus model and benchmarking against the current state of the art approach (11). Comp2GPR achieved high agreement with the reference model and substantially outperformed annotation-based methods, particularly in the reconstruction of multi-subunit enzyme complexes. Furthermore, integrating Comp2GPR-derived rules into the Yeast9 model leaded to improved gene essentiality predictions, yielding biologically meaningful corrections at the gene level while preserving overall model behavior. These results demonstrates that combining curated protein complex data with sequence-based validation provides a robust framework for improving GPR rule quality and enhancing the predictive performance of metabolic models.

## Methods

The GPR module provides an automated pipeline for annotating genome-scale metabolic models with Gene-Protein-Reaction (GPR) rules based on protein complex membership. Given a metabolic model (in SBML (14) or JSON format) containing the genes associated to each reaction based on functional annotation and a genome proteome (FASTA), the system identifies which genes encode subunits of known protein complexes and constructs Boolean GPR rules that capture obligate multi-subunit relationships. Where a single gene maps to multiple candidate complexes, the system flags the case for expert review, presenting detailed sequence similarity metrics to support informed disambiguation.

### GPR Database Construction

Protein complex information was sourced from the Complex Portal (EMBL-EBI) (2), a curated resource that aggregates protein complex data across multiple organisms. Raw data files were retrieved from the Complex Portal and included complex identifiers, component protein accessions, species annotations, functional descriptions, Gene Ontology (GO) terms (8), and supporting interaction evidence. To construct a gene–protein–reaction (GPR) database appropriate for genome-scale metabolic modeling, an AI-assisted curation workflow was applied to the raw complex data (16). The objective of this step was to retain only protein complexes relevant to metabolic models that explicitly represent metabolic reactions and their associated genes. Each protein complex was independently evaluated by an AI agent configured with domain-specific curation criteria reflecting standard assumptions in constraint-based metabolic modeling. The agent assessed the functional role of each complex using available annotations, including textual descriptions and GO terms. A complex was considered relevant only if it directly catalyzed a metabolic reaction or represented an obligate enzymatic assembly required for metabolic activity. Complexes whose annotated functions were restricted to non-metabolic processes—such as transcriptional regulation, signal transduction, chromatin organization, or other regulatory or structural roles—were excluded. The resulting curated dataset forms a GPR database optimized for metabolic model annotation, minimizing the inclusion of protein complexes that cannot be represented as metabolic reactions.

### Computational Workflow

The pipeline proceeds through five phases, executed as a background queued job on a persistent worker daemon.

#### Phase 1

Gene Matching. Genes associated with each reaction must be annotated in the metabolic model as starting point. The system extracts gene identifiers from the metabolic model and matches them against entries in the uploaded genome proteome FASTA file. Matching is bidirectional and flexible: gene ids are parsed from FASTA headers (e.g., tr|G0RHX3|G0RHX3 HYPJQ), SBML-style prefixes are stripped (e.g., G G0RHX3 corresponds to G0RHX3), gene names are extracted from GN= fields, and version suffixes (e.g., .1, .2) are removed to enable version-agnostic matching.

#### Phase 2

Complex Protein Discovery. Model gene sequences are searched against a pre-indexed reference database of characterized protein complex subunits using DIAMOND BLASTp (4), which provides 20–100*×* speedup over traditional BLAST (1). An e-value threshold of 1 *×* 10^*−*30^ is applied. To ensure deterministic results, a greedy 1-to-1 assignment algorithm resolves cases where multiple model genes match the same reference protein: genes are sorted by e-value, and each gene is assigned to the best-scoring unassigned identifier. For each assigned hit, all complexes containing that reference protein are retrieved from the complex database.

#### Phase 3

GPR Rule Construction. For each candidate complex, the system retrieves the full set of member proteins and their stoichiometric coefficients (format: P15790(2)|P25368(0)|P38930(1)|…), filtering out non-protein entries (e.g., CHEBI compound identifiers). Each member protein sequence is then searched against the genome using DIAMOND to identify the corresponding genome gene. The GPR rule is assembled as a Boolean AND of all matched genome genes, reflecting the obligate co-occurrence of subunits in a functional complex. Two quality filters are applied: (i) a coverage filter discards rules where fewer than 50% of complex members were matched (for complexes with more than three subunits), and (ii) a consistency filter discards rules where the original query gene is absent from the final rule, which would indicate that a different genome gene scored higher for the same complex member position.

#### Phase 4

Ambiguity Classification. Cases are categorized based on the number of candidate complexes per gene. A gene mapping to exactly one complex is classified as unique and auto-resolved.

A gene mapping to multiple complexes is classified as ambiguous and reserved for user review. An additional deduplication step auto-promotes ambiguous cases where all candidate complexes produce identical sets of genome genes in their GPR rules.

#### Phase 5

Rule Application and Merging. For unique cases, GPR rules are applied to all reactions containing the corresponding gene. When multiple complexes contribute rules to the same reaction, they are combined with Boolean OR. A merge strategy preserves non-overlapping branches from the original model GPR (containing only OR rules): the existing rule is split at top-level OR operators, branches with gene overlap are replaced by the new rule, and non-overlapping branches are retained. For example, if the original rule is A OR B and the new rule is (A AND C), the merged result is (A AND C) OR B.

### Ambiguity Resolution Interface

For ambiguous cases, a web-based review interface presents each case grouped by reaction. For each gene–complex pair, the interface displays the complex name, aliases, and biological description; the proposed GPR rule; the full list of complex member proteins with stoichiometry; and a BLAST mapping table showing genome gene matches with e-value, percent identity, and query coverage. The best hit is marked as selected, and alternative hits are available in a collapsible panel. Users may select one or more preferred complexes (combined with OR) before submitting, after which the same rule-building and merging logic is applied.

### Implementation

The module is implemented as a Django application within a larger metabolic modeling web platform. The data model comprises four entities: GPRAnalysisJob (job lifecycle and results), GPRBlastResult (cached DIAMOND search results), ComplexMetadata (complex descriptions for UI enrichment), and GPRAmbiguousCaseResolution (audit trail of user disambiguation decisions). Background processing is handled by a persistent worker daemon that polls the database at 5-second intervals to prevent race conditions in multi-worker deployments. Job status is communicated to the browser via AJAX polling at 2–3 second intervals. Access control is enforced per-session (anonymous), ensuring job privacy. A typical analysis completes in around one hour, dominated by the iterative DIAMOND searches for complex member proteins against the genome.

### Essentiality prediction

Experimentally validated essential genes for *Saccharomyces cerevisiae* were retrieved from the Saccharomyces Genome Database (SGD) (12). These annotations were used as the ground truth for evaluating model-based predictions of gene essentiality. A genome-scale metabolic model was used to predict essential genes through in silico gene knockout simulations. The yeast9 consensus model was loaded in Python using COBRApy (10), and gene–protein–reaction (GPR) associations were updated using externally derived GPR rules. Gene essentiality was evaluated using single gene deletion analysis implemented in COBRApy. Briefly, each gene in the model was individually knocked out by constraining the associated reactions according to the Boolean logic defined in the GPR rules. The resulting growth rate was computed using flux balance analysis (FBA) (18). Genes whose deletion resulted in zero predicted growth were classified as predicted essential, whereas all others were considered non-essential. To enable comparison with experimental data, all genes in the model were assigned binary labels: genes annotated as essential in SGD were labeled as non-viable (“N”), and the remaining genes were labeled as viable (“V”). The same labeling scheme was applied to the model predictions, where genes predicted to abolish growth were labeled as “N” and all others as “V”. Prediction performance was assessed using a normalized confusion matrix. The confusion matrix was computed using scikit-learn (17) and normalized by the total number of genes to allow comparison across methods. This representation summarizes the proportions of true positives, false positives, true negatives, and false negatives in the essentiality predictions.

## Results

### Protein Complex Database

Application of the AI-assisted curation workflow to the Complex Portal dataset resulted in a substantial refinement of the protein complex landscape toward metabolic relevance. The original dataset contained 5,295 protein complexes spanning a wide range of biological functions and organisms. Following systematic evaluation using domain-specific criteria, only 598 complexes were retained as directly relevant to metabolic modeling, corresponding to approximately 11% of the initial dataset. These curated complexes collectively comprise 1,558 unique protein components, forming the basis of a GPR-compatible protein complex database tailored for genome-scale metabolic model annotation. The large reduction in dataset size reflects the predominance of protein complexes in the Complex Portal that are associated with non-metabolic cellular processes, including transcriptional regulation, chromatin organization, signal transduction, and structural assembly. These complexes, while biologically important, cannot be directly mapped to metabolic reactions within the constraint-based modeling framework and therefore represent a source of potential noise when constructing gene–reaction associations. The filtering process effectively removes such entries by prioritizing complexes that either directly catalyze biochemical reactions or constitute obligate enzymatic assemblies required for metabolic function. The retained set of 598 complexes is enriched for functionally relevant enzymatic systems, including multimeric enzyme complexes and tightly coupled subunit assemblies that are essential for catalytic activity. By focusing on these metabolically actionable entities, the curated database ensures that the resulting GPR rules reflect biologically realistic constraints, such as subunit interdependence and the necessity of complete protein complexes for reaction activity. Importantly, the curated dataset reduces ambiguity in GPR construction by eliminating protein complexes with indirect or regulatory roles that could otherwise introduce incorrect logical relationships between genes and reactions. This is particularly relevant in cases where regulatory complexes, common ATP binding domains or protein–protein interaction assemblies are mistakenly interpreted as catalytic units, leading to erroneous AND relationships in GPR rules. By explicitly restricting the dataset to metabolically relevant complexes, the workflow minimizes such errors and improves the interpretability and mechanistic fidelity of the resulting gene–protein–reaction mappings. Overall, the resulting protein complex database provides a high-confidence foundation for GPR rule generation, balancing coverage of enzymatic diversity with strict relevance to metabolic function. The substantial reduction in dataset size, combined with the retention of key catalytic assemblies, demonstrates the effectiveness of the AI-assisted curation strategy in extracting actionable biochemical information from large, heterogeneous protein complex repositories.

### Comparison with GPRuler

I compared **Comp2GPR** with GPRuler(11), a tool designed for the automated reconstruction of gene–protein–reaction (GPR) rules from genomic and protein annotation data. GPRuler integrates information from resources such as UniProt (9), Metacyc (6) and Complex Portal (2) to infer both isozyme (OR) and multi-subunit enzyme (AND) relationships. While this strategy enables the reconstruction of complex enzyme associations, its reliance on annotation-derived mappings and database completeness can introduce ambiguity, including unsupported gene assignments and the incorporation of non-metabolic or spurious protein complexes.

In contrast, **Comp2GPR** reconstructs GPR rules directly from sequence-level evidence combined with curated protein complex information. The method enforces deterministic one-to-one gene– protein assignments and incorporates explicit filtering of spurious associations, resulting in more specific and reproducible GPR rules, particularly in cases where annotation coverage is incomplete or ambiguous.

These methodological differences were reflected in the agreement with the Yeast9 consensus GPR rules. After predicting the GPR rules of the Yeast9 consensus model (21), Comp2GPR achieved an overall exact-match accuracy of 95.9% (3962 exact matches out of 4131 evaluated rules), whereas GPRuler reached 31.3% accuracy (1293 exact matches out of 4131 evaluated rules). The difference was even more pronounced for reactions involving multi-subunit enzyme complexes represented by AND relationships. For the AND relationships present in Yeast9, Comp2GPR achieved 83.3% exact-match accuracy (174/209), while GPRuler correctly reconstructed only 1.4% of the rules (3/209). When considering the AND relationships present in the predicted rules, Comp2GPR recovered exact matches for 51.6% of the evaluated rules (174/337), whereas GPRuler achieved only 0.3% exact-match accuracy (3/891) (Figure 2). These results highlight the difficulty of inferring protein complexes from annotation alone and demonstrate the advantage of incorporating sequence-level validation, curated complex information, and deterministic assignment strategies for GPR rule reconstruction.

**Figure 1.**
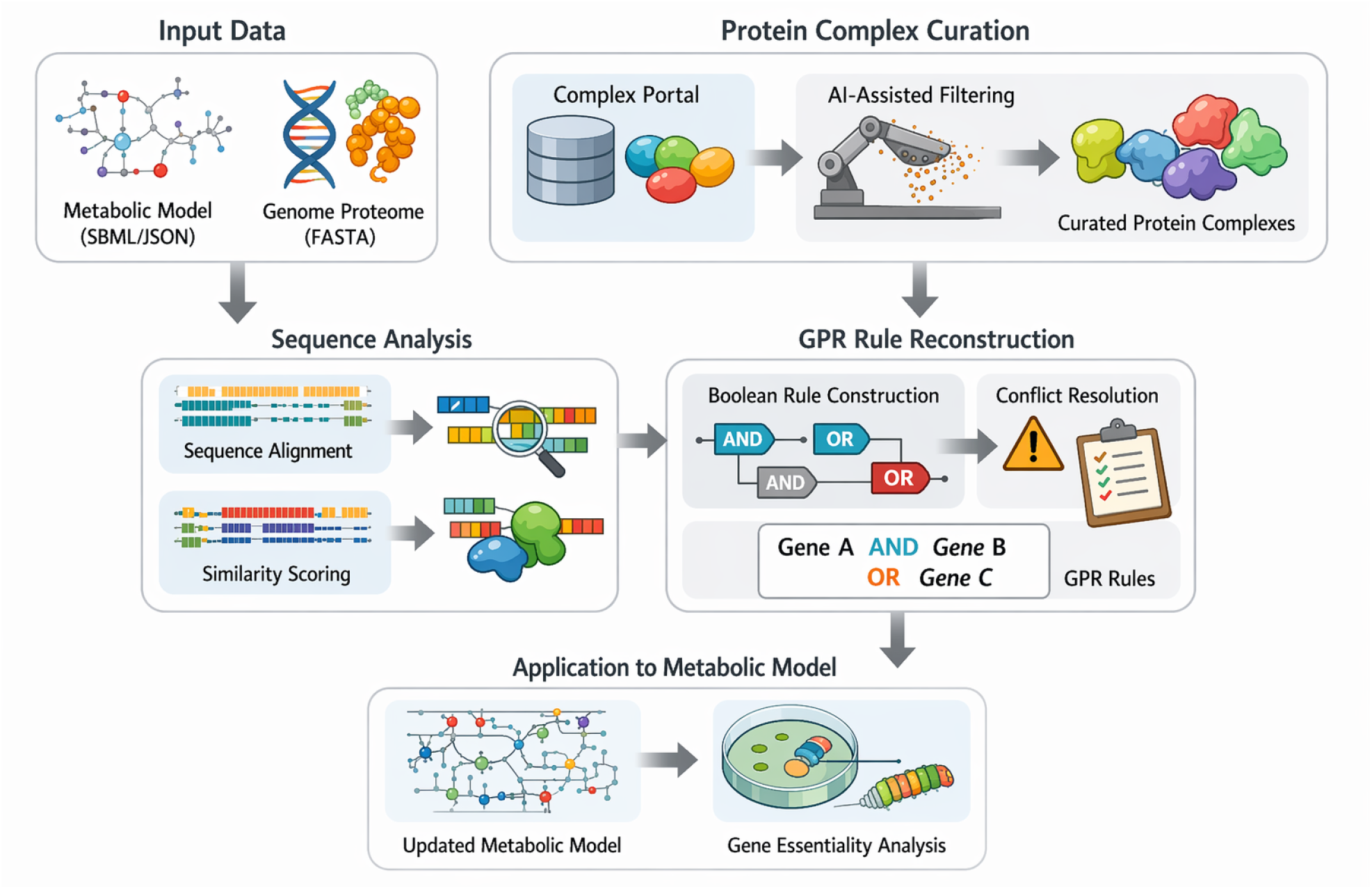
Comp2GPR pipeline for sequence-driven reconstruction of gene–protein–reaction (GPR) rules. The workflow integrates a genome-scale metabolic model, a proteome (FASTA), and a AI curated protein complex database. Model genes are matched to protein sequences and searched against reference complex subunits using sequence similarity, followed by Boolean rule construction capturing multi-subunit (AND) and isoenzyme (OR) relationships. Ambiguous assignments are classified for resolution, and validated rules are applied and merged into the model, generating an updated representation of gene–reaction associations.

**Figure 2.**
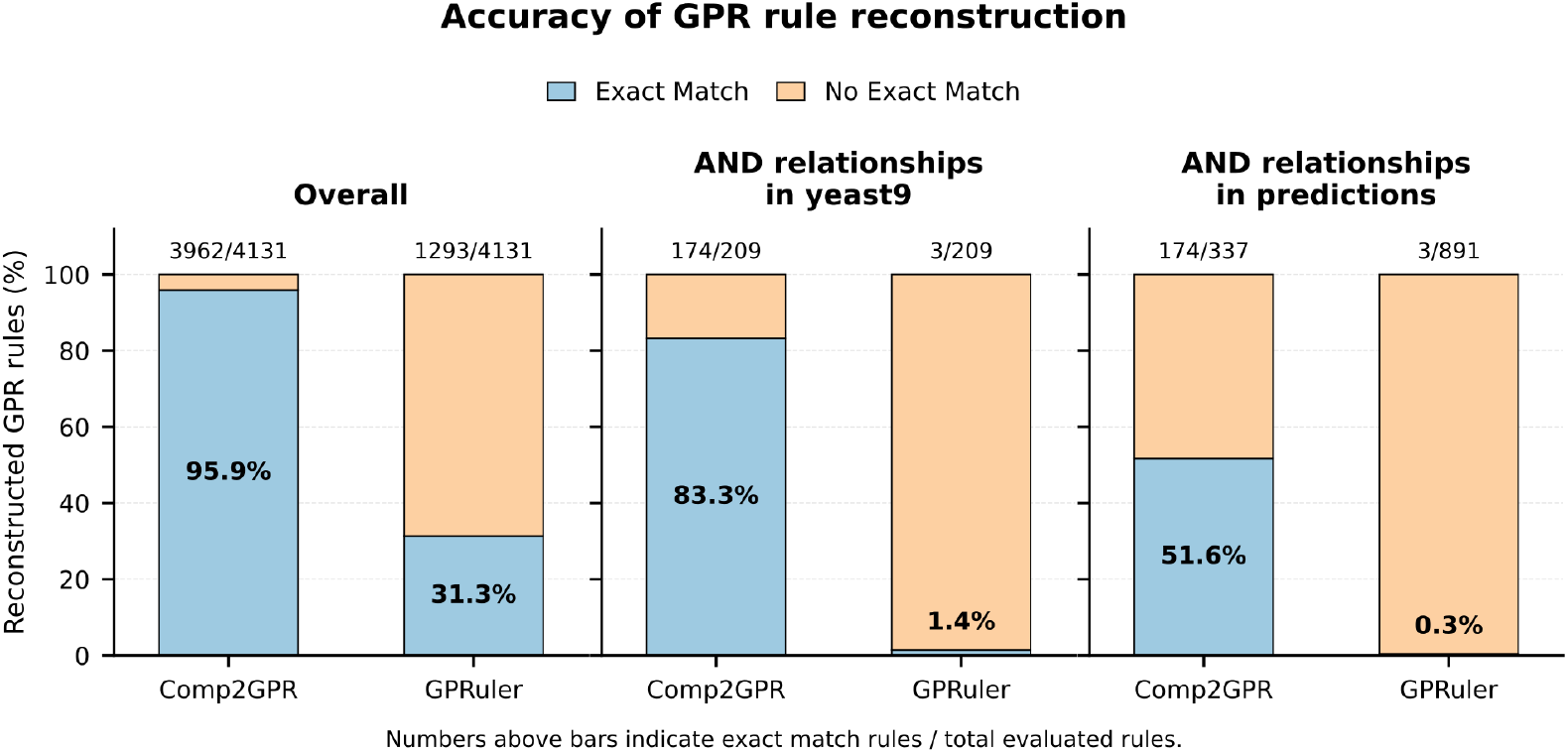
Comparison of exact-match accuracy for GPR rule reconstruction of Yeast9 consensus model. Stacked bars show the percentage of exact and non-exact matches obtained by Comp2GPR and GPRuler for all evaluated reactions, AND relationships in yeast9, and AND relationships in the predicted rules. Numbers above bars indicate exact matches over total evaluated rules. Comp2GPR showed higher exact-match accuracy across all subsets, especially for AND relationships.

### Comparison with Yeast9 consensus model

To assess the functional impact of GPR rules generated by **Comp2GPR**, I compared gene essentiality predictions obtained with the original Yeast9 model to those obtained after replacing the original GPR rules with Comp2GPR- and GPRuler-derived rules (Figure 3A–C).

**Figure 3.**
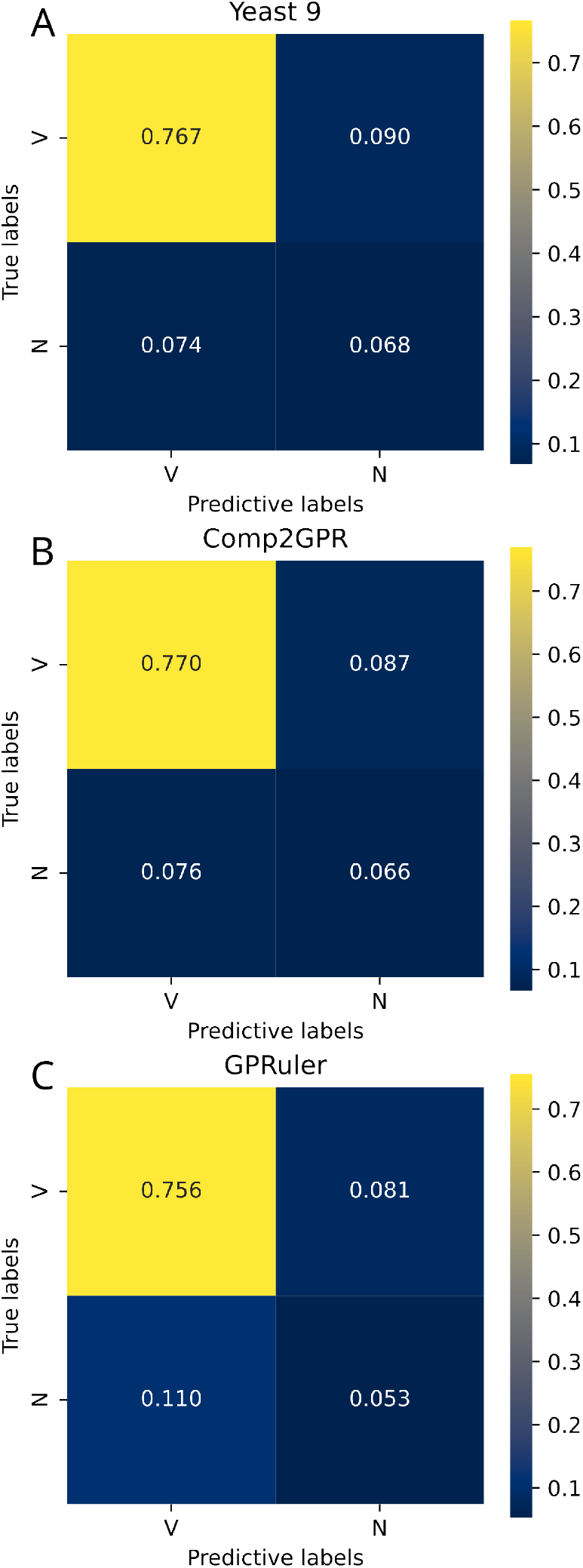
Comparison of gene essentiality predictions using different GPR rule sets.(A) Original Yeast9 model.(B) Yeast9 model updated with Comp2GPR-derived rules.(C) Yeast9 model updated with GPRuler-derived rules. Each panel shows the confusion matrix comparing predicted versus experimental gene essentiality, where values represent fractions of genes in each category. V indicates genes experimentally classified as non-essential (viable), and N indicates essential genes (Non-viable).

Overall, updating the model with Comp2GPR rules preserved global predictive behavior. The confusion matrices for essential versus non-essential gene predictions (Figure 3A–B) show highly similar distributions. The fraction of experimentally viable genes correctly predicted as non-essential increased slightly from 0.767 in Yeast9 to 0.770 with Comp2GPR, while the fraction of non-essential genes incorrectly predicted as essential decreased from 0.090 to 0.087. The fraction of essential genes incorrectly predicted as non-essential increased marginally (from 0.074 to 0.076), whereas the prediction of essential genes slightly decreased (0.068 vs. 0.066). These small shifts indicate that Comp2GPR refines gene–reaction associations without destabilizing overall model behavior.

In contrast, GPRuler-derived rules led to a noticeable degradation in predictive performance (Figure 3C). The fraction of essential genes incorrectly predicted as non-essential increased to 0.110, while the fraction of correctly predicted viable genes decreased to 0.756. Although the misclassification of non-essential genes as essential slightly decreased (0.081), this was accompanied by a reduced ability to capture essential genes accurately.

Despite the modest aggregate changes observed with Comp2GPR, gene-level analysis revealed substantial qualitative improvements. The updated model correctly reclassified 21 genes as non-essential that were previously predicted as essential or absent from the original Yeast9 predictions (YGL037C, YJR025C, YJR073C, YKR072C, YOR054C, YPR167C, YNL138W, YGR102C, YLR188W, YMR096W, YJL062W, YOR239W, YER145C, YKL190W, YOR101W, YCR094W, YDR221W, YNR048W, YNL323W, YER072W, and YPL019C). These corrections are consistent with known limitations in GPR reconstruction, where missing isoenzymes, incomplete protein complexes, or overly restrictive AND relationships can artificially introduce essential bottlenecks.

In addition, Comp2GPR rescued two experimentally essential genes (YGR277C and YMR301C) that were incorrectly predicted as non-essential in the original model. These cases likely reflect improved representation of protein complex requirements, preventing biologically implausible flux bypasses.

These improvements were achieved with limited trade-offs. Only three genes (YBR029C, YER043C, and YHR007C) were correctly classified as essential in the original model but misclassified after updating, and three misclassified genes (YCL009C, YFR015C, and YJL071W) showed the opposite behavior.

Overall, replacing Yeast9 GPR rules with those generated by Comp2GPR improves biological consistency and introduces localized corrections in gene essentiality predictions while maintaining stable global model performance. These results indicate that the benefits of Comp2GPR arise from mechanistic improvements in gene–reaction logic rather than from global constraint relaxation.

## Discussion

Accurate reconstruction of gene–protein–reaction (GPR) rules remains a critical challenge in genome-scale metabolic modeling, particularly for reactions involving multi-subunit enzyme complexes. The results presented in this study demonstrate that **Comp2GPR** substantially improves the reconstruction of GPR rules compared to annotation-driven approaches, with especially strong gains in cases requiring AND relationships.

The high agreement of Comp2GPR with the Yeast9 consensus model (95.9% accuracy) indicates that combining curated protein complex information with sequence-level validation is an effective strategy for reconstructing GPR rules. In contrast, the markedly lower accuracy of GPRuler (29%) highlights the limitations of relying primarily on annotation-based mappings. These differences become even more pronounced when considering multi-subunit enzyme complexes, where Comp2GPR achieves over 83.3% accuracy, while GPRuler fails almost entirely (1.4%). This result underscores the inherent difficulty of accurately inferring obligate subunit relationships from annotation data alone and confirms that explicit sequence-based validation is essential for resolving these associations.

The performance gap observed in AND-rule reconstruction is particularly relevant for metabolic modeling. Errors in multi-subunit enzyme representation can introduce severe functional inconsistencies, such as artificially blocking reactions when required subunits are missing or, conversely, allowing biologically implausible flux when complexes are incompletely specified. The improved performance of Comp2GPR in this context reflects its ability to enforce mechanistically consistent gene–protein mappings, ensuring that all required subunits are simultaneously represented. This deterministic and constraint-aware strategy contrasts with annotation-driven approaches, which may introduce ambiguous or over-permissive rules due to incomplete or noisy database information.

Despite the large differences in GPR reconstruction accuracy, the impact on global gene essentiality predictions is relatively modest. This observation suggests that genome-scale models exhibit a degree of robustness to local inaccuracies in GPR rules, likely due to redundancy in metabolic networks and the presence of alternative pathways. However, gene-level analysis reveals that improvements in GPR quality translate into biologically meaningful corrections, including the recovery of previously misclassified non-essential genes and the restoration of essentiality for genes that were absent or incorrectly predicted as non-essential. These corrections are consistent with improved representation of isoenzyme redundancy and protein complex requirements, indicating that Comp2GPR enhances the mechanistic fidelity of gene–reaction associations rather than simply relaxing or tightening constraints globally.

A key strength of Comp2GPR lies in its integration of curated protein complex data with a deterministic sequence-based mapping strategy. By explicitly filtering non-metabolic complexes and enforcing one-to-one gene–protein assignments, the method reduces ambiguity and improves reproducibility. The substantial reduction of the Complex Portal dataset to a metabolically relevant subset further contributes to the robustness of the approach by minimizing the inclusion of irrelevant or misleading complexes. Together, these design choices address two major sources of error in GPR reconstruction: database noise and ambiguous gene mappings.

Nevertheless, several limitations should be considered. First, the approach relies on the availability and quality of curated protein complex data, which may be incomplete for less well-characterized organisms. Second, gene-to-reaction annotations are required. Although these annotations do not need to be complete—since the pipeline systematically searches for all components of each complex—their absence may result in missing complex information. Third, while the method includes an ambiguity resolution pipeline, manual intervention may still be necessary in complex cases involving multiple plausible mappings.

Future work could address these limitations by expanding the curated complex database, incorporating probabilistic or confidence-weighted assignments for ambiguous cases, and integrating complementary data types such as structural predictions or co-expression evidence.

In conclusion, this study demonstrates that accurate modeling of protein complexes is a key determinant of GPR rule quality and that sequence-driven reconstruction strategies provide a substantial advantage over annotation-based methods. By improving the representation of gene–protein relationships, Comp2GPR enhances the biological realism of metabolic models and supports more reliable simulation of genotype–phenotype relationships.

## Conclusion

In this study, I introduced **Comp2GPR**, an automated framework for reconstructing gene–protein–reaction (GPR) rules by integrating curated protein complex information with sequence-level validation. The method achieves high agreement with the Yeast9 consensus model and substantially outperforms annotation-based approaches, particularly in the reconstruction of multi-subunit enzyme complexes.

Our results demonstrate that accurate modeling of protein complexes represents a key bottleneck in GPR reconstruction. By enforcing deterministic gene–protein assignments and filtering non-metabolic or unsupported complexes, Comp2GPR improves the mechanistic fidelity and reproducibility of GPR rules. These improvements are especially pronounced for AND relationships, where annotation-driven methods struggle to correctly represent obligate subunit dependencies.

Importantly, integrating Comp2GPR-derived rules into a genome-scale metabolic model leads to biologically meaningful corrections in gene essentiality predictions while preserving overall model behavior. This indicates that improvements in GPR rule quality translate into more accurate representation of genotype–phenotype relationships without compromising model robustness.

Overall, Comp2GPR provides a robust and scalable framework for enhancing metabolic model annotations. By combining curated complex knowledge with sequence-based validation, the method advances the reconstruction of gene–reaction associations and supports more reliable simulation and analysis of cellular metabolism.

## Abbreviations

GPR: 
GEM: 
GO: 
BLAST: 
FASTA: 
SBML: 
AI: 

## Conflicts of interest

The authors declare that they have no competing interests.

## Funding

This study was funded by FoodID (2025-2027), an NSF Global Centers project supported by Business Finland (3545/31/2024), Research Council of Finland (366905) and by FinBioFAB (2025-2029), an NSF Global Centers project supported by Business Finland (3479/31/2024) and the Research Council of Finland (365981).

## Data availability

Comp2GPR is available as a standalone package via the GitHub repository https://github.com/vttresearch/Comp2GPR and can also be accessed through a web interface at https://carvefungi.vtt.fi.

## Notes

https://carvefungi.vtt.fi

### Competing Interest Statement

The authors have declared no competing interest.

